# Overexpression of *MusaVicilin* Gene for Disease Resistance in Banana

**DOI:** 10.1101/2025.10.01.679877

**Authors:** Sarah Wanjiku Macharia, Jaindra Nath Tripathi, Valentine Otang Ntui, Samwel Muiruri Kariuki, Leena Tripathi

## Abstract

Banana Xanthomonas wilt (BXW) disease, caused by *Xanthomonas vasicola* pv. *musacearum*, is a major constraint to banana production in East and Central Africa. All cultivated banana varieties are susceptible, with the wild progenitor *Musa balbisiana* being the only known source of complete resistance. Limitations in classical breeding have prompted the exploration of molecular genetic tools, such as genetic modification, to develop resistant cultivars. Comparative transcriptomic analyses revealed a five-fold upregulation of *MusaVicilin* gene in *M. balbisiana* (BB genome) compared to the BXW- susceptible ‘Pisang Awak’ when challenged with the pathogen, suggesting its role in defense. This study investigated whether constitutive overexpression of the *MusaVicilin* gene cloned from *M. balbisiana* could enhance resistance to BXW in the susceptible ‘Sukali Ndiizi’ cultivar (AAB genome). Transgenic events were developed with the *MusaVicilin* gene under the control of the constitutive CaMV 35S promoter. These events exhibited enhanced disease resistance compared with non-transgenic control plants. The overexpression of *MusaVicilin* highlights its potential as a candidate gene for engineering resistance to BXW in susceptible cultivars. Moreover, *MusaVicilin* could serve as a valuable component in gene stacking strategies aimed at developing durable, disease-resistant banana varieties.

## Introduction

Bananas (*Musa* spp.) hold critical importance in global food security, feeding over 400 million people worldwide^1^. Cultivated in more than 140 countries in the subtropics and tropics, bananas have a global annual production of approximately 183.6 million tonnes, with India as the leading producer at approximately 36.6 million tonnes^2^. Bananas are among the most widely consumed fruits globally, with consumption projected to reach 90 million metric tonnes by 2026^3^. In East Africa, bananas contribute significantly to daily per-person calorie intake, ranging from 30% to 60%, with Uganda having the highest consumption rates^4^. Cultivated bananas are allopolyploids or autopolyploids of *Musa acuminata* (AA genome) and *Musa balbisiana* (BB genome). The diversity of banana cultivars is estimated to be 300 to 1,200, belonging to various genomic groups such as AA, BB, AB, AAA, AAB, ABB, AAAA, AAAB, AABB, and ABBB^5^.

Banana production in East and Central Africa faces severe challenges due to Banana Xanthomonas Wilt (BXW), a systemic bacterial disease caused by *Xanthomonas vasicola* pv. *musacearum (Xvm)*^6^. This devastating disease affects all cultivated banana and can lead to yield losses of up to 100 %^7^. BXW has resulted in the destruction of entire plantations in affected regions, compelling farmers to seek infection-free planting sites^8^. The economic toll of BXW is estimated to range from $2 to 8 billion over the course of a decade^9^. Transmission of *Xvm* occurs through various means, including insect vectors, contaminated tools, soil-borne bacterial inoculum, and infected planting material^7^.

In the absence of BXW-resistant cultivated varieties, a comprehensive approach is required for effective management of BXW disease. Strategies such as rigorous phytosanitary measures, careful decapitation of male buds^10^, and strict adherence to hygiene protocols^7^ are crucial for preventing disease transmission. However, such strategies are inherently limited: their effectiveness depends heavily on farmer compliance, they are labor-intensive, and they cannot fully eliminate the pathogen, which may also adapt to management practices^11^. This emphasizes the need for development of improved banana varieties with enhanced resistance to BXW, in addition to continued research into comprehensive control startegies^10, 12, 13^.

While all cultivated banana varieties are vulnerable to BXW disease, the wild progenitor *M. balbisiana* exhibits complete resistance^7, 14^. Moreover, Nakato et al.^12^ noted significant *Xvm* tolerance in the AA genome of Zebrina species, implying the existence of tolerant traits within the current banana germplasm. Transferring the disease resistance trait from wild-type banana *M. balbisiana* to farmer-preferred cultivars through conventional breeding is a lengthy and challenging process due to the sterility^15^ of most cultivars coupled with polyploidy and the long generation times. In addition, the presence of banana streak virus (BSV) sequences in the B genome restricts the utilization of *M. balbisiana* in conventional breeding^16^. These barriers highlight the urgency for innovative biotechnological interventions such as genetic engineering to revolutionize banana breeding, ultimately fostering the development of resilient and improved banana varieties.

Genetic engineering has shown promising progress by incorporating defense genes such as *Hrap, Pflp, Xa21*, *ES-Pflp,* and *AtEFR* into banana plants^13, 14, 15, 16, 17^. However, the continual evolution of new pathogenic strains and the adaptability of *Xvm* pose ongoing challenges to disease eradication efforts. To overcome these obstacles, it is crucial to explore the genetic potential of the BXW-resistant wild banana varieties^18, 19, 20, 14^.

Comparative transcriptomics between BXW-susceptible banana cultivar ‘Pisang Awak’ and BXW-resistant *M. balbisiana* during early *Xvm* infection identified numerous defense-related genes. Notably, *MusaVicilin* was strongly upregulated in *M. balbisiana* but downregulated in the susceptible cultivar, highlighting the rationale for overexpressing *MusaVicilin* in susceptible banana cultivars to enhance resistance^14^.

Vicilins are 7S globulins with molecular weights typically between 200 and 400 kDa and exhibit a complex range of functions. They are predominant in legumes and are primarily known as seed storage proteins. They also serve as precursors for antimicrobial peptides (AMPs)^21^, specifically vicilin-like AMPs, that are released through proteolytic cleavage during seed development or in response to pathogen attack. During this process, vicilin proteins are cleaved at precise sites by vacuolar processing enzymes, or defense-related proteases, releasing smaller bioactive vicilin-like peptides that exhibit antimicrobial properties^21, 22, 23^. These AMPs form an integral part of the plant’s innate immune system, combating bacteria, fungi, viruses, yeast and pests ^24–36^.

Vicilin-like peptides have demonstrated significant antimicrobial properties, contributing to plant defense against various pathogens. Vicilin-like proteins from *Capsicum baccatum* L., narrow- leafed lupin, and *Clitoria fairchildiana* exhibited antifungal and insecticidal properties^30, 31^. Cystein-rich vicilin peptides from *Centrosema virginianum* improved fungal resistance in transgenic tobacco^32^, while antimicrobial peptides derived from the N-terminus region of vicilin proteins in *Macadamia integrifolia* and *Theobroma cacao*^33^, also displayed potent antifungal activity against *Fusarium oxysporum* and *Aschochyta rabiae.* Additionally, Almeida et al.^34^ revealed antibacterial activity in hydrolysates of cowpea vicilin. Further, binding to chitin, vicilins from cowpea, and *Enterolobium contortisiliquum* seeds interfered with insect larval development and prevented fungal germination^35, 36^. These findings collectively highlight the broad-spectrum defensive capabilities of vicilin-derived antimicrobial peptides against bacterial, fungal, and insect pests across diverse plant species.

Building on these insights and previous transcriptome analysis, *MusaVicilin* was identified as a candidate resistance gene in BXW-resistant *M. balbisiana*. To assess its functional role, this study constitutively overexpresses *MusaVicilin* in the BXW-susceptible cultivar ‘Sukali Ndiizi’. Transgenic events overexpressing *MusaVicilin* were evaluated under greenhouse conditions against *Xvm* challenge. The results showed markedly enhanced resistance compared with non-transgenic controls, confirming that *MusaVicilin* contributes to BXW defense and represents a promising target for developing resistant banana cultivars.

## Materials and methods

### Plant material

Embryogenic cell suspension (ECS) of banana cultivar ‘Sukali Ndiizi’ used for transformation were generated from immature male flowers^37^, and maintained at 28±2°C on a rotary shaker at 95 rpm in the dark.

### *MusaVicilin* gene construct design

The *MusaVicilin* gene sequence was downloaded from banana genome B (*M.balbisiana*), gene ID: Vicilin ITC1587_Bchr3_P08153 (Mba01_g02390.1) from the Banana Genome Hub^38^. PCR primers (Mb_Vicilin_F: AGAGCGAGCGAGAGAGAGAA and Mb_Vicilin_R: CCTCTTCCCTTCACTCTGT) were designed to isolate the full-length gene (1664 bp) from *M. balbisiana*. PCR amplification was done using Hotstar Taq DNA polymerase (QIAGEN, Germany). After amplification, the PCR product was resolved in 1% agarose gel stained with GelRed (Biotium, San Francisco, USA), a fluorescent nucleic acid stain. The PCR product was purified (using PCR purification kit (QIAGEN, Hilden, Germany), according to manufacturers’ instructions) and cloned into the Gateway entry vector, pCR8/GW/TOPO (Invitrogen, Carlsbad, CA, USA). The ligated product was transformed into *DH5α E. coli* competent cells and selected on LB medium containing spectinomycin (100 mg/l). Ten colonies were selected, and plasmid DNA was extracted. It was initially verified by PCR and later by Sanger sequencing using M13 primers. The transformants containing the correct *M. balbisiana* gene sequences were subcloned in the sense orientation between the attR1 and attR2 recombination sites in the binary vector pMDC32 by LR clonase^TM^ (Invitrogen, Carlsbad, CA, USA) recombination reaction to yield pMDC32*_MbVicilin*. The plasmid was transformed into *DH5α E. coli* competent cells and selected on LB medium containing kanamycin (50 mg/l). Ten independent colonies were selected and verified by digestion with *Eco*RI. The colony with the correct insert was introduced into the highly virulent *Agrobacterium tumefaciens* strain EHA105 by electroporation. Ten colonies were selected, and colony PCR was performed to check integration of the plasmid into the *Agrobacterium*. The pMDC32 vector contained *hygromycin phosphotransferase (hpt)* as the selection marker gene, and the *MusaVicilin* gene was driven by the cauliflower mosaic virus (CaMV) 35S promoter.

### Generation of transgenic plants

Transgenic events overexpressing *MusaVicilin* were generated by delivering the pMDC32- 35S::*MbVicilin* plasmid into the ECS of ‘Sukali Ndiizi’ by *Agrobacterium*-mediated transformation as described previously by Tripathi et al.^37^. The *Agrobacterium*-infected ECS were regenerated on a selective medium supplemented with 12.5 mg/l hygromycin following the protocol described by Tripathi et al.^37^. All regenerated putative transgenic events were maintained and multiplied on proliferation medium at 28±2°C under a 16/8h light/dark photoperiod with fluorescent lighting. The putative transgenic events were validated for the presence of the *hpt* gene by PCR. The transgenic shoots were transferred to rooting medium and the well-rooted plants were subsequently transferred to soil in the greenhouse for BXW resistance evaluation.

### Molecular characterization of transgenic events

#### PCR analysis

Genomic DNA was extracted from freshly collected leaf samples of the putatively transformed events and non-transgenic control (NTC) plants using the cetyltrimethylammonium bromide (CTAB) method^39^. All hygromycin-resistant (65 events) were assessed for transgenesis by targeting a 415 bp long region of the *hpt* gene using the primers *hpt* forward (5′ GATGTTGGCGACCTCGT 3′ and *hpt* reverse 5′ GTGTCACGTTGCAAGACCTG 3’). The PCR reaction was set up in a 25-μL reaction volume containing 2.5μL 10X PCR buffer, 0.3μL 10mM dNTPs, 1μL of 10μM reverse and forward primers, 2μL genomic DNA (100ng/μL), 0.2μL Taq DNA polymerase of 5 units/μL (Qiagen, Germany), and 18μL nuclease-free water. BIO-RAD T100 Thermo Cycler conditions were as follows: initial denaturation at 95°C for 5 min, followed by 32 cycles of denaturation at 94°C for 30 s, annealing at 56°C for 30 s, and extension at 72°C for 1 min, then final extension at 72°C for 7 min. Genomic DNA from NTC plants was used as a negative control, and the pMDC32-35S::*MbVicilin* plasmid DNA was used as a positive control. The amplified products were separated on a 1% agarose gel (Duchefa, Netherlands) stained with GelRed (Biotium, San Francisco, USA) and visualized under ultraviolet light using Syngene™ Ingenius 3 gel documentation system.

### Southern blot analysis

Southern blot analysis was performed on the *MusaVicilin* overexpression events as per the method described by Tripathi et al.^10^. Briefly, 10μg of genomic DNA from each sample was digested with the restriction enzyme *Hind*III at 37° C for 12 h and separated on 0.8% agarose gel at 50 V for 5h. The plasmid pMDC32-35S::*MbVicilin* and genomic DNA from a NTC plant were used as positive and negative controls, respectively. The gel, stained with GelRed (Biotium, San Francisco, USA), was viewed under ultraviolet light (using Syngene™ Ingenius 3 gel documentation system) to confirm the digestion. The restricted DNA was denatured, then blotted onto a positively charged membrane (Roche Diagnostics, West Sussex, UK) and fixed by cross- linking in ultraviolet light (UV 500 Crosslinker, Amersham Biosciences). The membrane was then hybridized with a 415-bp *hpt-*specific probe, PCR-labeled with digoxigenin (DIG). Hybridization and probe detection were performed using a DIG Luminescent Detection Kit for Nucleic Acids (Roche Diagnostics, UK) per the manufacturer’s protocol.

### Plant growth analysis

A subset of 11 PCR-positive transgenic events and NTC plants were analyzed for growth parameters in the greenhouse. Three replicates of well-rooted plants for each event were transferred to soil in small plastic cups (10 cm diameter) and acclimatized for 30 days in a humidity chamber, then transferred to bigger pots (30 cm diameter) and grown in the greenhouse for 90 days at 25–30°C. The growth parameter data, including plant height, pseudostem girth, number of functional leaves, and length and width of the middle leaf, were recorded from 90- day-old plants. The total leaf area was calculated using the formula below^40^;

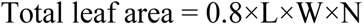

In which L = length of the middle leaf, W = width of the middle leaf, and N = total number of leaves in the plant.

### Disease evaluation under greenhouse conditions

Transgenic events, and NTC plants were evaluated for resistance against *Xvm* under greenhouse conditions. Each transgenic event and NTC plant were replicated three times. A total of 52 events were used for the disease assay. These transgenic events and NTC plants were arranged in a completely randomized design. The Ugandan strain of *Xvm* used for disease screening was sourced from a -80° C maintained culture in the International Institute of Tropical Agriculture (IITA), Nairobi laboratory. The *Xvm* inoculum was cultured in plates in YPGA medium (0.5% yeast extract, 0.5% peptone, 1% glucose, and 0.8% micro agar) for 48 h at 28°C as previously described by Mwangi et al.^41^. A single colony was aseptically isolated and cultured for 48 h in 50 ml of YPG medium at 28°C in an incubator shaker (200 rpm). Subsequently, the liquid culture was centrifuged at 4000 rpm for 15 min, and the pellet was resuspended in sterile distilled water. The bacterial culture was adjusted to an OD_600nm_ of 1 using sterile distilled water. The second open functional leaf of 90-day-old potted plant was inoculated with 100μL of the bacterial culture using an insulin syringe. The plants were maintained in the greenhouse under observation, and symptoms were recorded over a 60-day post-inoculation (dpi) period. The data collected included the number of days for appearance of the first symptom, number of days for complete wilting, disease severity, and the number of leaves showing symptoms at 60 dpi. The data was used to calculate percent resistance compared to NTC plants using the formula below^15^;

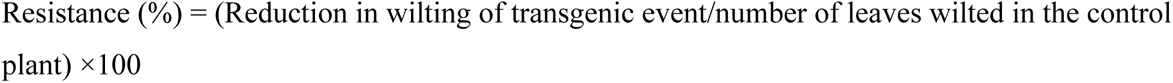

In which, the reduction in wilting was the total number of leaves minus the number of leaves wilted. Disease severity was rated on a scale of 0–5 (0- no signs of the disease, 1- a single leaf showing symptoms, 2-two to three leaves with symptoms, 3- four to five leaves with symptoms, 4- all the leaves have symptoms but the plant not dead, 5- complete death).

Plants were categorized as resistant if they did not show any symptoms, partial resistant if the symptoms did not spread to all the leaves, and susceptible if the symptoms spread to all the leaves, causing complete wilting or death of the plant.

### Gene expression analysis

A subset of 11transgenic events overexpressing *MusaVicilin* and NTC plants were selected based on their disease evaluation results in the greenhouse. To determine the relative gene expression of these events compared to the control NTC plant, total RNA was extracted from the leaves of two-week-old shoots using the Qiagen Plant RNeasy Kit, adhering to the manufacturer’s instructions. The RNA quality and concentration were determined using NanoDrop 2000 (Thermo Fisher Scientific). Subsequently, 1μg of total RNA was converted to cDNA using the LunaScript^®^ RT SuperMix cDNA Synthesis Kit (New England BioLabs, E3010L) as outlined in the user manual. The resulting cDNA was diluted ten-fold, and 5μl was used as template for qRT-PCR with primers (*MusaVicilin* forward 5′ TCAGGGCGAGTCGATAATA 3′ and *MusaVicilin* reverse 5′ CTCTTCTTGGCTTCCTCTTC) and SYBR Green Master Mix (Applied Biosystems) on a QuantStudio5 real-time PCR system (Applied Biosystems, Thermo Fisher Scientific). The Musa 25S forward 5′ ACATTGTCAGGTGGGGAGTT 3′ and Musa 25S reverse primer: 5′ CCTTTTG TTCCACACGAGATT 3′ served as the internal control^26^. A non-template control, containing nuclease-free water, was also included in the analysis. Each sample consisted of three technical replicates. The reaction volume was 20μl, consisting of 10μl of 5X Luna Universal qPCR Master Mix, 0.2μl of 10μM forward and reverse primers, 5μl cDNA, and 4.6μl nuclease-free water. Real-time PCR conditions were as follows: initial denaturation at 95°C for 5 min, followed by 40 cycles of denaturation at 94°C for 30 s, annealing at 60°C for 30 s, and extension at 72°C for 1 min. The relative gene expression levels were determined using the 2−ΔCt method^42^.

### Correlation analysis of transgenic events

Correlation analyses were conducted to investigate relationships among disease resistance percentages, relative gene expression (fold change), and transgene copy number across 11 *MusaVicilin* transgenic events and NTC plants. The relative gene expression levels were quantified using qRT-PCR and normalized relative to the NTC^42^. Transgene copy number was determined using Southern blot analysis. Disease resistance was assessed as a percentage as previously described by Tripathi et al.^15^. Pearson’s correlation analysis was conducted to determine the linear relationships among the three variables using the cor() function in R software (version 4.4.3). The statistical significance of the pairwise correlations was evaluated using the cor.test() function, and a matrix of p-values was generated using the cor_pmat() function. The results were visualized in a matrix format using ggcorrplot. Prior to analysis, data distributions were examined for normality to justify the use of Pearson correlation. Statistical significance was evaluated at p ≤ 0.05. Each transgenic event and NTC were treated as independent data points (n=12). Correlations were interpreted to identify potential associations between transgene expression, copy number, and resistance phenotype.

### Statistical analysis

Minitab Statistical Software, version 17 (Pennsylvania, USA), was used to analyze the data. One-way analysis of variance (ANOVA) was used to compare the differences in disease resistance and plant growth traits across different transgenic events and NTC plants. Fisher’s HSD test was used to determine the significant difference between the means, and statistical significance was determined at p ≤ 0.05. Correlation analyses between gene expression levels and phenotypic traits were performed using R software (version 4.4.3)

## Results

### *MusaVicilin* plasmid construct

The *MusaVicilin* gene (ITC1587_Bchr3_P08153; Mba01_g02390.1) was successfully amplified from *M. balbisiana* using PCR primers (Mb_Vicilin_F/R), yielding a 1,664 bp amplicon. Cloning into the Gateway entry vector pCR8/GW/TOPO and subsequent LR recombination into the binary vector pMDC32 generated the plasmid construct pMDC32*_MbVicilin* (Figure 1A), verified by colony PCR, *EcoR*I digestion, and Sanger sequencing. The construct, harboring the *MusaVicilin* gene under the *CaMV* 35S promoter and hygromycin selection marker (Figure 1B), was confirmed in *Agrobacterium tumefaciens* EHA105 via colony PCR.

**Figure 1:**
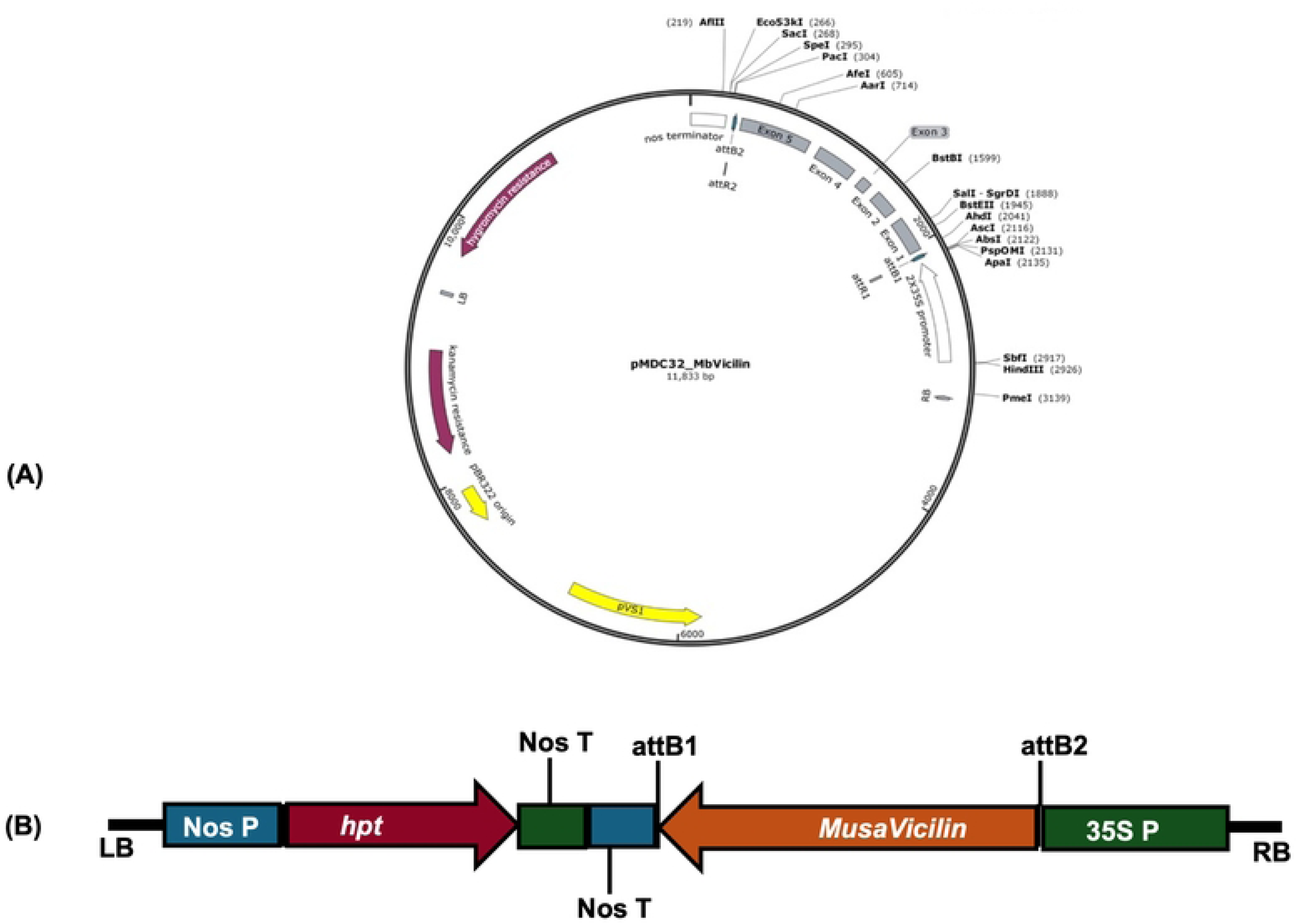
Schematic presentation of *MusaVicilin* plasmid construct. (A) Plasmid structure of pMDC32_*MbVicilin*; (B) *MusaVicilin* gene expression cassette as structured in the pMDC32_*MbVicilin* vector, LB; left border, Nos P; Nopaline Synthase Promoter, *hpt; hygromycin phosphotransferase* selectable marker gene, Nos T; Nopaline Synthase Terminator, attB1; attachment site B1, aatB2; attachment site B2, 35S P; Cauliflower Mosaic Virus (CaMV) 35S promoter, RB right border.

### Generation of transgenic events

‘Sukali Ndiizi’ ECSs were successfully transformed with *Agrobacterium tumefaciens* strain EHA105 containing the pMDC32-35S::*MbVicilin* plasmid. The transformed cells were positively selected on hygromycin 12.5 mg/L containing medium, whereas untransformed cells turned black (Figure 2A). A total of 65 independent events were generated. Transgenic events were multiplied (Figure 2B-D), then successfully developed roots (Figure 2E) within 3–4 weeks across all independent events. Following acclimatization, well-rooted plantlets of the transgenic events were established in the greenhouse for growth analysis and disease evaluation (Figure 2F). These events were maintained in proliferation medium (PM) under 16 h/8 h photoperiod at 26–28 °C, subculturing every 6–8 weeks.

**Figure 2:**
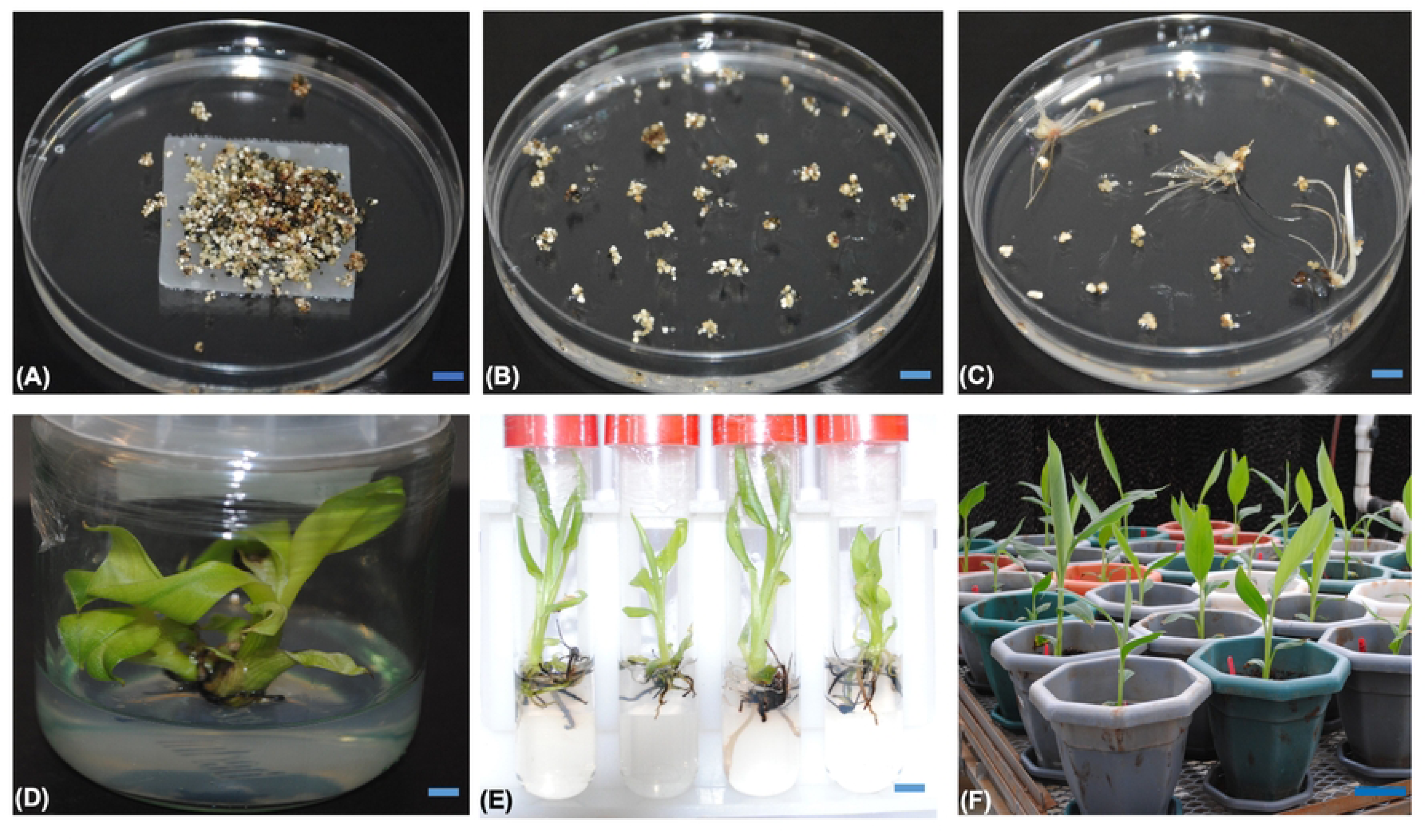
Generation of banana transgenic events. (A) Embryogenic cells in selective embryo development medium; (B) Embryos maturing on selective embryo maturation medium; (C) Germination of embryos in selective embryo germination medium; (D) Multiplication of plantlets in proliferation medium; (E) Well-rooted plantlets in proliferation medium (PM); (F) Potted events in the greenhouse awaiting disease assay. The scale bar represents 1 cm for panels (A) – (E), and 10 cm for panel (F).

### Molecular characterization of transgenic events PCR analysis

The regenerated putative *MusaVicilin* transgenic events were validated for the presence of the *hpt* transgene using PCR with *hpt*-specific primers. The tested events showed the expected 415 bp amplicon, confirming successful transgenesis (Figure 3A). No amplification was observed in the NTC plants.

**Figure 3:**
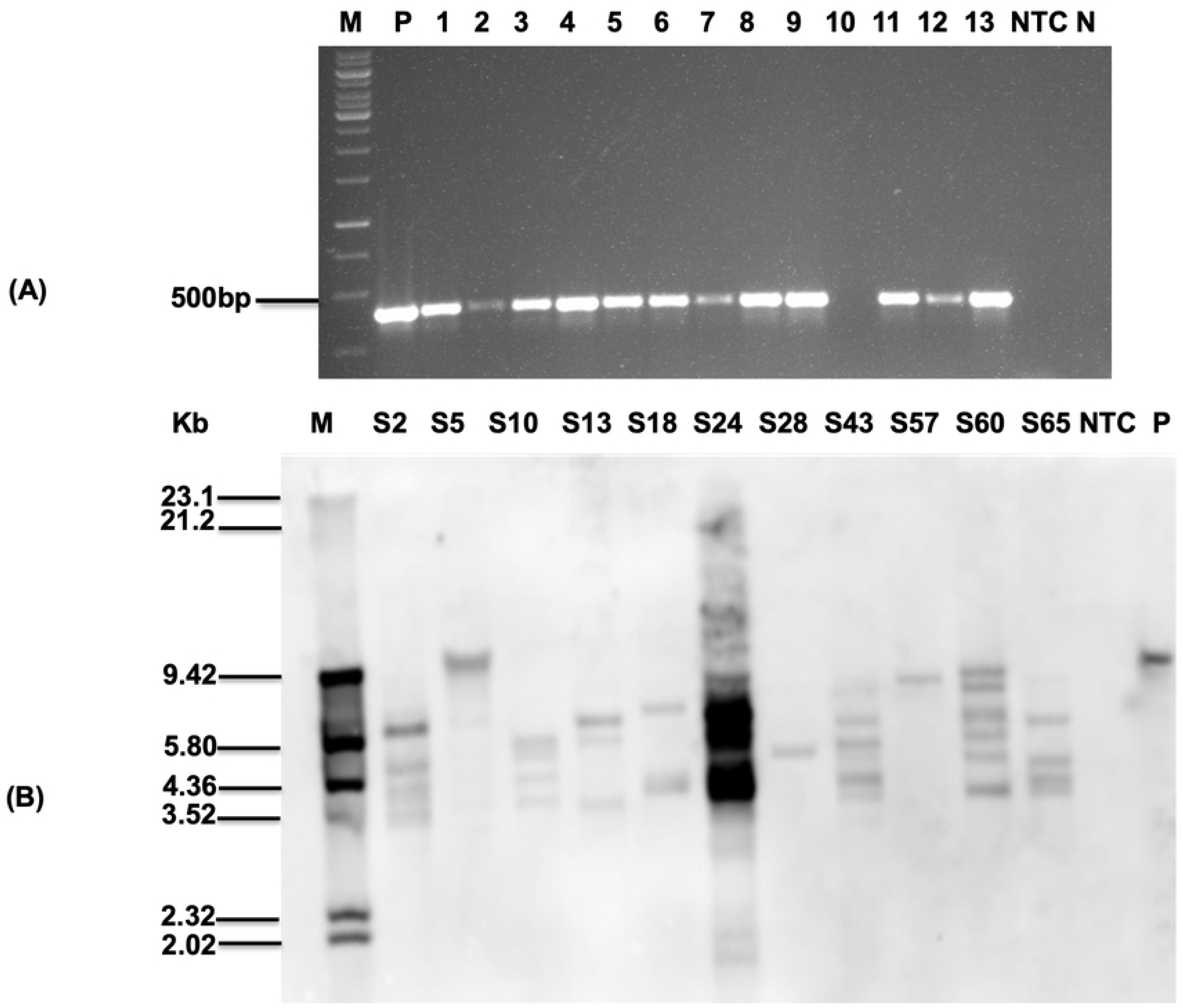
Molecular analysis of putative transgenic banana events. (A) Representative polymerase chain reaction assay to confirm the presence of *hpt* gene, M; 1kb DNA ladder, 1-13; transgenic events, P; Plasmid, NTC; Non-transgenic control, N, non-template control; (B) Southern blot analysis to confirm the integration of *MusaVicilin* gene, M; molecular marker, NTC; Non-transgenic control, P; plasmid.

### Southern blot analysis

To verify transgene integration and copy number, the Southern blot technique was employed to evaluate 11 PCR-positive events (S2, S5, S10, S13, S18, S24, S28, S43, S57, S60, S65) from *pMDC32-35S::MbVicilin*. Southern blot hybridization of *Hind*III-digested genomic DNA using *hpt* probe confirmed the integration of the transgene in the plant genome with different hybridization profiles, indicating random insertion of the transgene in the genome of the tested events. The transgene copy number incorporated in the different events ranged from one to multiple (Figure 3B). Events S5, S28, and S57 had a single gene copy, with events S13 and S18 having three gene copies, while event S60 had the most gene copies (6). No transgene integration was detected in the NTC plant.

### Plant growth analysis of transgenic events

To evaluate the impact of overexpression of *MusaVicilin* gene on plant morphology, 11 events (S2, S5, S10, S13, S18, S24, S28, S43, S57, S60, S65), and NTC plants were selected, and the following growth parameters measured: plant height, pseudostem girth, total leaf area, and number of functioning leaves. No morphological defects were detected, and all the *MusaVicilin* events grew normally. Also, no significant differences (p ≤ 0.05) were observed in the different growth parameters between the transgenic events and the NTC plants, suggesting that overexpression of the *MusaVicilin* gene in the transgenic events did not cause any adverse growth effect (Figure 4).

**Figure 4:**
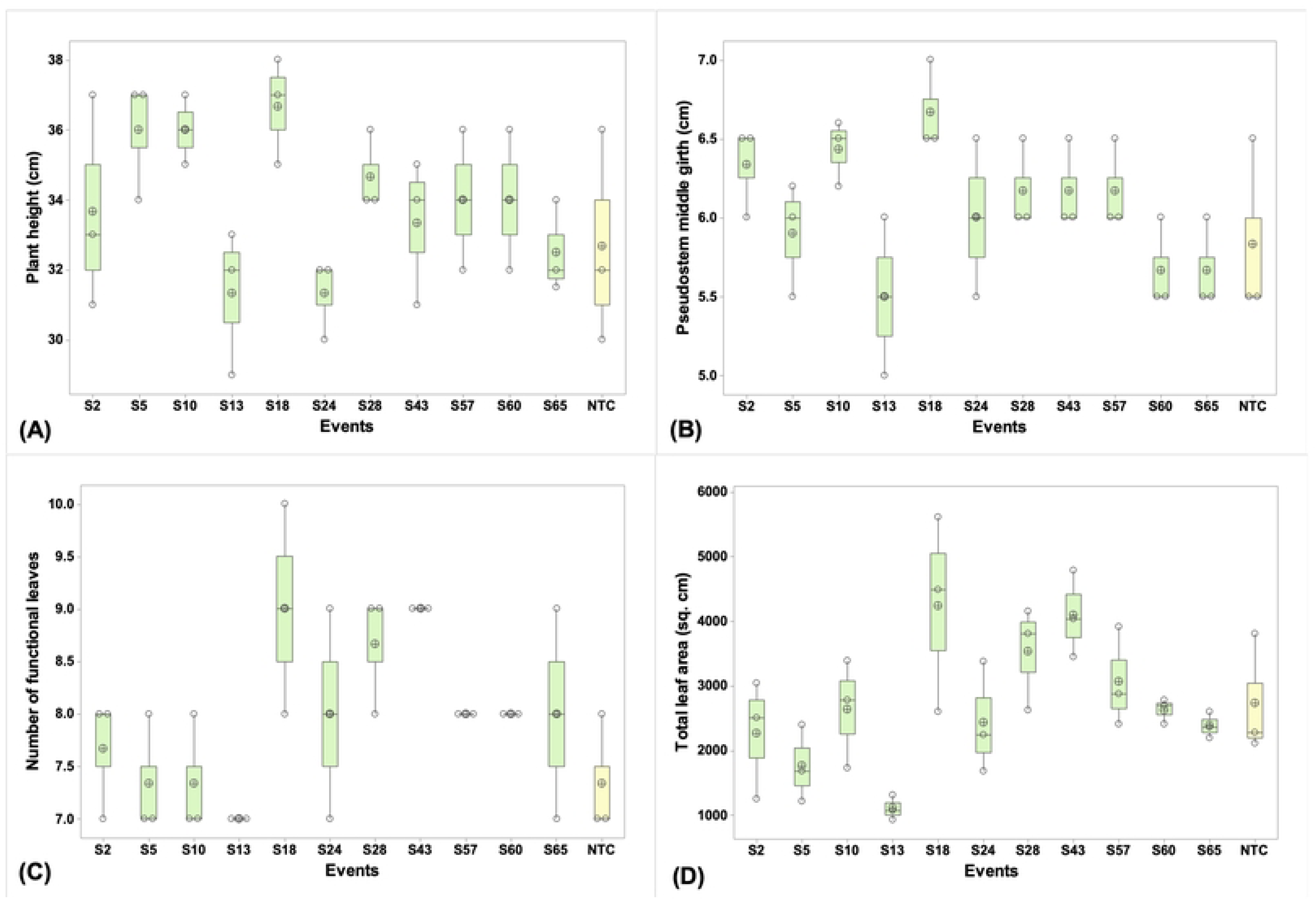
Growth analysis of banana transgenic events. (A) Plant height, B) Pseudostem middle girth, (C) Number of functional leaves, (D) Total leaf area. Data are presented as means ± standard deviation. Bars with different letters are significantly different at p ≤ 0.05 according to Fisher’s HSD test (n = 3).

### Evaluation of transgenic events for disease resistance

To assess the response of transgenic events to the bacterial pathogen *Xvm*, 52 events, along with NTC plants, underwent evaluation for disease resistance under greenhouse conditions. The NTC plants exhibited pronounced symptoms (chlorosis or necrosis) after about 16 dpi, which spread throughout the entire plant, culminating to complete wilting at 22.67±4.93 dpi (Table 1) with a 100% disease incidence (Figure 5C). This confirmed their susceptibility to *Xvm* infection. In contrast, event S2 exhibited complete resistance against *Xvm* (Figure 5A), displaying no disease symptoms upon *Xvm* challenge (Table 1). Additionally, event S5 (Figure 5B) demonstrated a substantial disease resistance (95.2%), with the manifestation of disease symptoms limited solely to the inoculated leaf with a very low disease severity index of 4.8%. The remaining 50 events showed partial resistance against *Xvm*, ranging from 25% to 77.8%, with only about 15% of the events showing susceptibility comparable to that of the NTC plants. This comprehensively unveils a spectrum of disease resistance levels among the *MusaVicilin* events, signifying the efficacy of constitutive overexpression of the *MusaVicilin* gene in conferring protection against *Xvm* infection.

**Figure 5:**
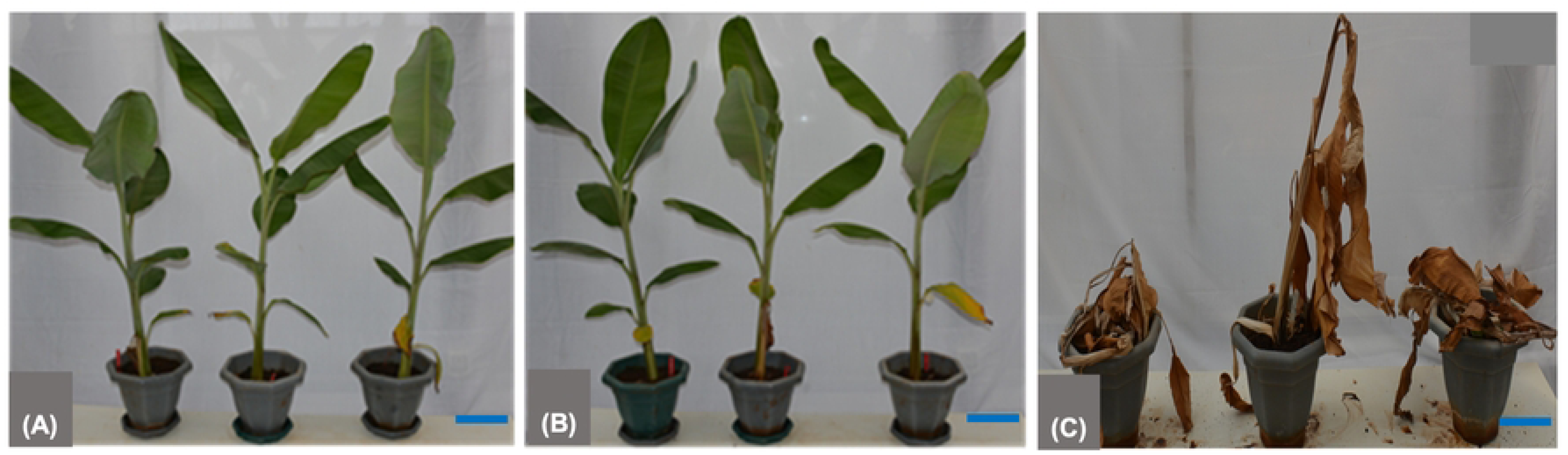
Disease evaluation of banana transgenic events against *Xanthomonas vasicola* pv. *musacearum*. (A) Transgenic banana event S2 showing complete resistance to BXW, (B) Event S57 showing wilting symptom solely on the inoculated leaf of the second replicate plant, (C) Non-transgenic control exhibiting complete wilting. Plants were assessed for 60 days post inoculation with *Xvm*. The scale bar represents 10 cm.

**Table 1.**
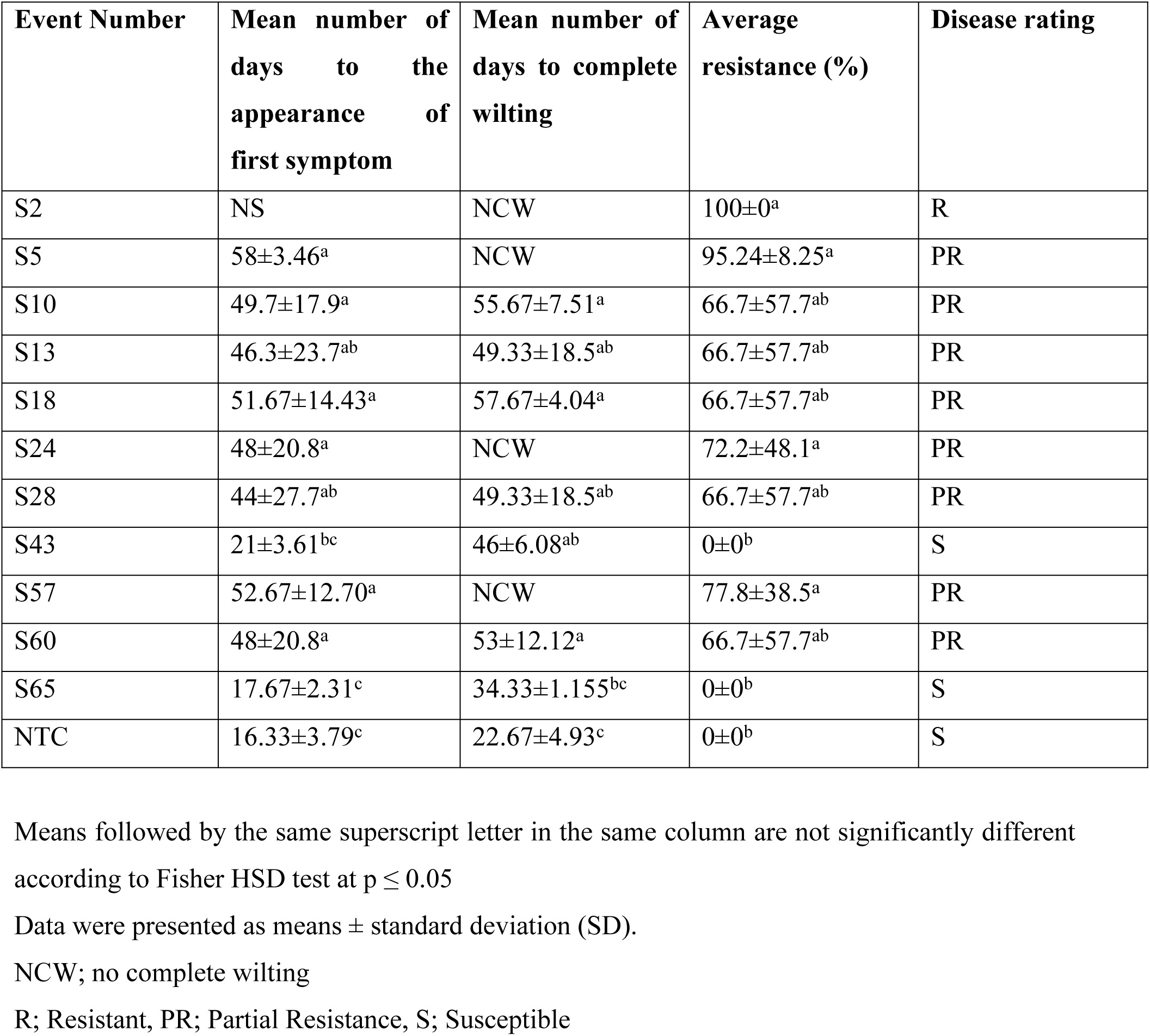
Greenhouse evaluation of transgenic events for resistance against *Xanthomonas vasicola* pv. *musacearum*.

| Event Number | Mean number of days to the appearance of first symptom | Mean number of days to complete wilting | Average resistance (%) | Disease rating |
| --- | --- | --- | --- | --- |
| S2 | NS | NCW | 100±0<sup>a</sup> | R |
| S5 | 58±3.46<sup>a</sup> | NCW | 95.24±8.25<sup>a</sup> | PR |
| S10 | 49.7±17.9<sup>a</sup> | 55.67±7.51<sup>a</sup> | 66.7±57.7<sup>ab</sup> | PR |
| S13 | 46.3±23.7<sup>ab</sup> | 49.33±18.5<sup>ab</sup> | 66.7±57.7<sup>ab</sup> | PR |
| S18 | 51.67±14.43<sup>a</sup> | 57.67±4.04<sup>a</sup> | 66.7±57.7<sup>ab</sup> | PR |
| S24 | 48±20.8<sup>a</sup> | NCW | 72.2±48.1<sup>a</sup> | PR |
| S28 | 44±27.7<sup>ab</sup> | 49.33±18.5<sup>ab</sup> | 66.7±57.7<sup>ab</sup> | PR |
| S43 | 21±3.61<sup>bc</sup> | 46±6.08<sup>ab</sup> | 0±0<sup>b</sup> | S |
| S57 | 52.67±12.70<sup>a</sup> | NCW | 77.8±38.5<sup>a</sup> | PR |
| S60 | 48±20.8<sup>a</sup> | 53±12.12<sup>a</sup> | 66.7±57.7<sup>ab</sup> | PR |
| S65 | 17.67±2.31<sup>c</sup> | 34.33±1.155<sup>bc</sup> | 0±0<sup>b</sup> | S |
| NTC | 16.33±3.79<sup>c</sup> | 22.67±4.93<sup>c</sup> | 0±0<sup>b</sup> | S |

Means followed by the same superscript letter in the same column are not significantly different according to Fisher HSD test at $p \leq 0.05$
Data were presented as means ± standard deviation (SD).
NCW; no complete wilting
R; Resistant, PR; Partial Resistance, S; Susceptible

### Relative expression of the *MusaVicilin* gene

Eleven transgenic events (S2, S5, S10, S13, S18, S24, S28, S43, S57, S60, S65), representing a range of BXW resistance phenotypes observed in greenhouse screening, were selected for qRT-PCR analysis of the *MusaVicilin* gene. The selected events represented a range of tolerance levels from 0% (non-transgenic control) to 100% (Table 1). Differences in *MusaVicilin* transcript levels were observed between the transgenic events investigated compared to NTC. For instance, event S57 exhibited the highest *MusaVicilin* gene expression level (over 1350-fold), followed by events S13 and S5. Event S43 exhibited the least fold of relative expression (120-fold) (Figure 6). These results confirm successful overexpression of the *MusaVicilin* gene in the transgenic events.

**Figure 6:**
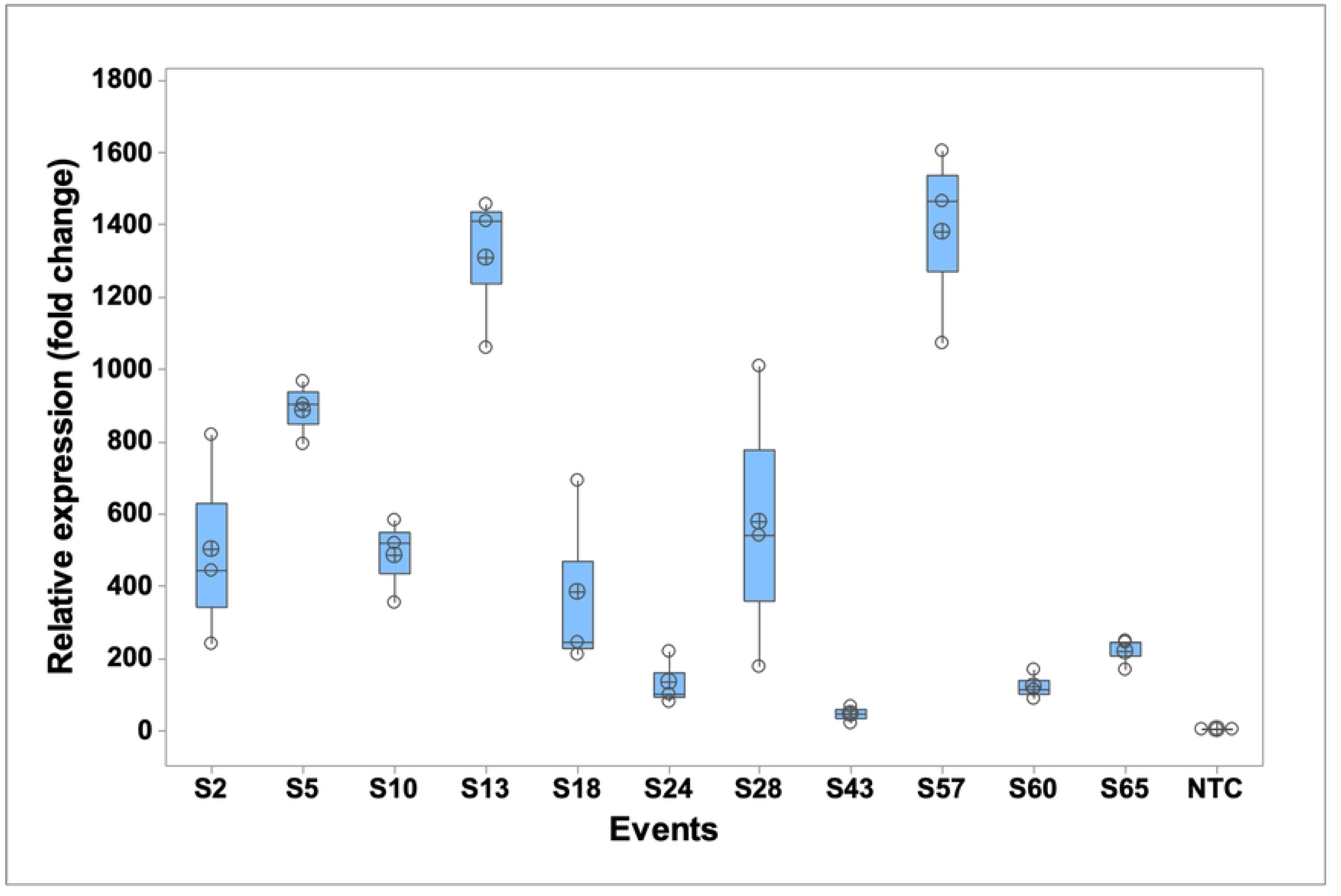
*MusaVicilin* relative expression in transgenic events. qRT-PCR analysis of constitutive overexpression of *MusaVicilin* events.

### Correlation analysis of transgenic events

Correlation analysis among disease resistance percentages, relative gene expression (fold change), and transgene copy number revealed distinct patterns across the 11 transgenic events (S2, S5, S10, S13, S18, S24, S28, S43, S57, S60, S65) and NTC plant. A moderate positive correlation was observed between gene expression fold change and resistance percentages (r = 0.53) (Figure 7A), suggesting that higher transgene expression may contribute to enhanced disease resistance. However, this correlation did not reach statistical significance (p = 0.08) (Figure 7B). In contrast, the relationship between transgene copy number and disease resistance percentages was weak and non-significant (r = 0.07, p = 0.82), indicating that variation in copy number did not have a discernible impact on resistance levels. This suggests that having more copies of the transgene does not necessarily translate into increased resistance. Interestingly, a negative correlation was observed between transgene copy number and gene expression (r = - 0.35), although this was also not statistically significant (p = 0.27).

**Figure 7:**
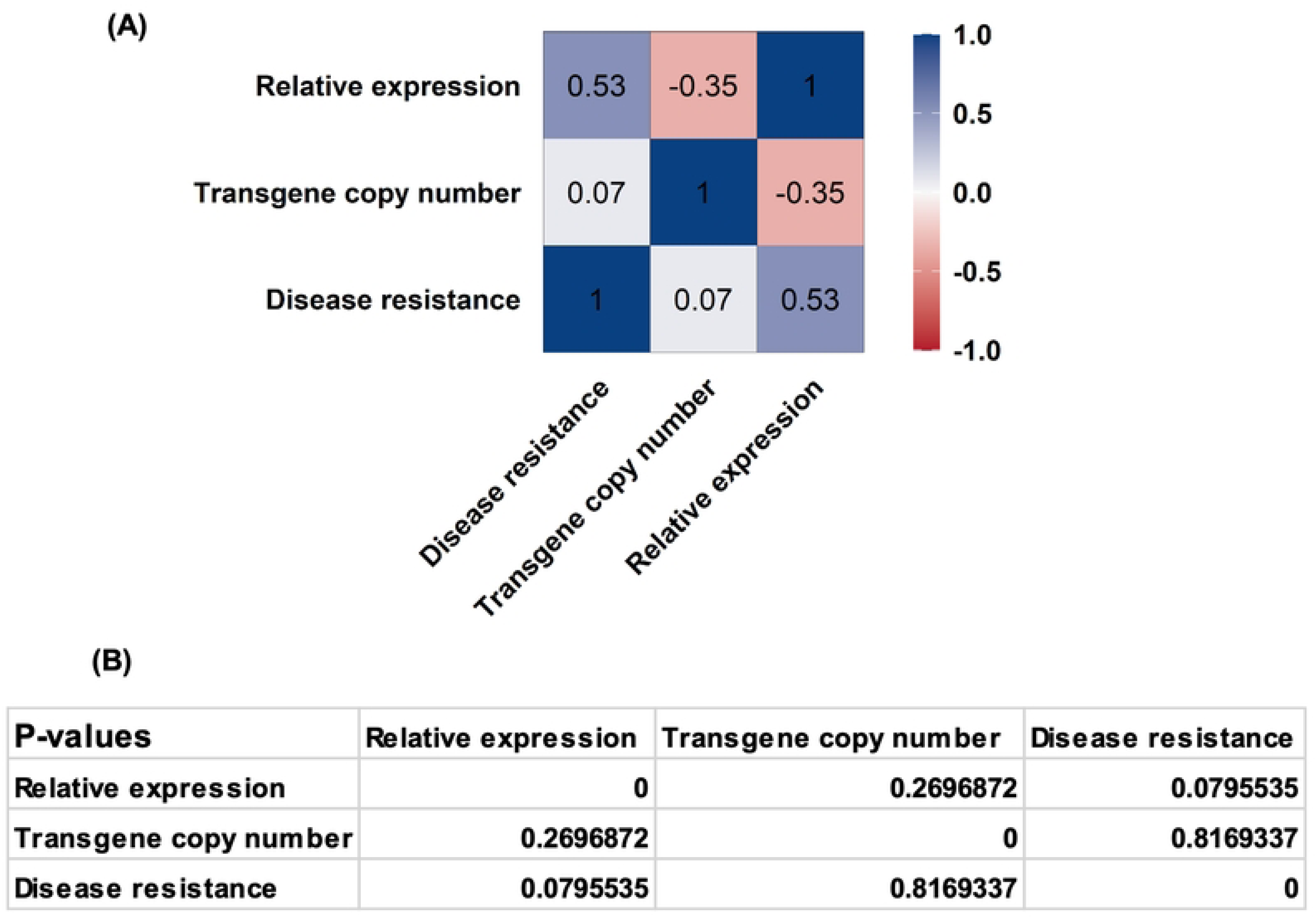
Pairwise Pearson correlation analysis of transgenic events. (A) Matrix showing Pearson correlation coefficients (r values) for all pairwise comparisons between disease resistance percentages, transgene copy number, and relative gene expression (fold change) across the transgenic events (11 events) and a non-transgenic control (NTC). The r values indicate the strength and direction of linear relationships between each variable pair, with values closer to +1 or -1 representing stronger positive or negative correlations, respectively, and values near 0 indicating weak or no correlation, (B) Table showing the corresponding p-values for each pairwise Pearson correlation in (A), indicating the statistical significance of the observed correlations. No statistically significant correlations were found among these variables (all p ˃ 0.05).

## DISCUSSION

BXW poses a significant threat to banana production in East and central Africa, impacting millions of livelihoods and regional food security. Conventional management strategies, including cultural practices and breeding for resistance, have had limited success in controlling BXW due to the genetic complexity of banana breeding and the limited availability of resistant traits^43^. The absence of fertile diploid bananas completely resistant to BXW further limits traditional breeding efforts^7, 15, 17^. To overcome these challenges, modern biotechnological approaches such as genetic transformation offer promising alternatives by introducing and overexpressing resistance genes to develop BXW-resistant banana varieties.

Antimicrobial peptides (AMPs) are crucial plant defense molecules with broad activity against pathogens^24, 25, 26^. Vicilin-like proteins release potent AMPs that provide effective protection against various disease-causing pathogens and pests^27, 28, 29^. The upregulation of the *MusaVicilin* gene, a vicilin-like AMP during BXW infection in resistance to *Musa balbisiana* at early infection, further supports its role in plant defense^21^. By overexpressing the *MusaVicilin* gene in the BXW susceptible banana, the colonization of *Xvm* could be restricted. We confirmed this hypothesis by constitutively overexpressing the *MusaVicilin* gene in susceptible banana cultivar ‘Sukali Ndiizi’, and the transgenic events generated were evaluated for their response against *Xvm* under greenhouse conditions. These assays revealed that transgenic banana overexpressing *MusaVicilin* exhibited enhanced resistance to BXW disease compared to NTC plants.

The *MusaVicilin* transgenic events demonstrated a spectrum of resistance against *Xvm* infection, underscoring its potential in bacterial wilt defense. The complete resistance observed in event S2, coupled with the near complete resistance in event S5 (95.2%), highlights the capacity of the *MusaVicilin* gene to block pathogen progression when stably expressed at optimal levels effectively. This aligns with studies in *Capsicum baccatum L., Clitoria fairchildiana, Macadamia integrifolia, Theobroma cacao, Centrosema virginianum, Enterolobium contortisiliquum,* and cowpea, where vicilin-like AMPs disrupted bacterial cell membranes or suppressed virulence factors, thereby curtailing disease spread^30–36^. Partial resistance (25% - 75%) observed in the remaining events suggested that transgene expression levels, copy number, or insertion sites critically influence gene efficacy, consistent with findings in other AMP-overexpression systems^26^. The non-transgenic control plants wilted rapidly (22.67 ± 4.93dpi), compared to the resistant events that delayed pathogen colonization, comparable with other vicilin-like AMPs^23^. Though 15% events remained susceptible, most showed improved disease resistance over NTC plants, confirming the potential of the *MusaVicilin* gene in bacterial wilt management. The observed variation in resistance levels among the transgenic events likely reflects positional effects of transgene insertion, which can significantly influence gene expression and stability^44^. Integration near heterochromatic regions, transcriptionally inactive zones, or regulatory elements may lead to variable or silenced expression, thereby impacting the resistance phenotype. Hence, thorough molecular characterization and screening of multiple independent transformation events are essential in understanding and managing positional effects, ensuring the successful development of stable and effective transgenic disease-resistant plants^45, 46^.

The *MusaVicilin* transcript levels in the transgenic events varied widely and did not consistently correlate with transgene copy number. For instance, event S2, with four gene copies, exhibited approximately 400-fold overexpression, achieving complete resistance to *Xvm*, whereas event S60, despite having six gene copies, showed only a 220-fold overexpression and no resistance. Similarly, events S57 (77.8%) and S13 (66.7%) demonstrated extremely high expression levels (up to 1,300-fold) with just one and three gene copies, respectively, while event S24, carrying five gene copies, had a comparatively low 120-fold expression. Event S43, with four copies, showed a mere 4-fold increase in transcript levels and succumbed to the disease. This confirms the moderate positive correlation between gene expression and disease resistance observed (r = 0.53, p = 0.08), suggesting that increased transgene expression may enhance resistance, although this relationship was not statistically significant. In contrast, transgene copy number showed no significant correlation with resistance (r = 0.07, p = 0.82) and was weakly negatively correlated with gene expression (r = –0.35, p = 0.27). Such discrepancies suggest that transgene copy number alone does not determine expression or resistance. Instead, factors such as positional effects of transgene insertion^44^, promoter activity^47^, transcriptional silencing^48^, and epigenetic regulation^49^ likely play critical roles in modulating expression levels. Moreover, the threshold of *MusaVicilin* expression required for effective defense against BXW may not be linearly related to copy number but rather to the efficiency of transcript accumulation and subsequent protein activity. These findings highlight key points especially successful resistance requires achieving sufficiently high *MusaVicilin* expression, regardless of copy number, and future work should focus on optimizing expression stability, possibly through targeted transgene integration, promoter engineering, or genome-editing approaches.

Overexpression of the *MusaVicilin* gene in the ’Sukali Ndiizi’ banana cultivar, did not induce noticeable adverse effects on plant morphology under greenhouse conditions. Transgenic events were phenotypically indistinguishable from NTC plants, suggesting that *MusaVicilin* overexpression may not negatively impact yield, aligning with findings in other crops like cucumber, cotton, brassica, and soybean^50, 51, 52, 53^. However, these results were obtained under controlled greenhouse conditions, and confined field trials are necessary for a definitive assessment of the gene’s impact on agronomic traits and overall productivity under realistic growing conditions.

Antimicrobial peptides (AMPs) slow disease progression by disrupting pathogen cell membranes, inhibiting enzyme activities, and interfering with pathogen replication^54^. These mechanisms reduce pathogen load and slow down the spread of infection within the plant. AMPs contribute to both basal and non-host disease resistances by providing a broad-spectrum defense against a wide range of pathogens^55^. This dual role is reflected in their ability to enhance quantitative resistance by delaying symptom appearance and reducing disease severity, as well as providing qualitative resistance by entirely preventing pathogen infection in some cases^56, 57, 58^.

In this study, the overexpression of the *MusaVicilin* gene in the transgenic events demonstrated quantitative resistance to BXW. Specifically, these transgenic events showed delayed disease symptom onset ranging up to 58 days, compared to 12 days for NTC plants. Furthermore, the transgenic events exhibited a lower disease severity (38.3%), compared to 100% for NTC plants. These findings demonstrate that *MusaVicilin* overexpression enhances quantitative resistance to BXW, providing a promising approach for banana disease management.

In summary, this study demonstrates that transgenic banana overexpressing the *MusaVicilin* antimicrobial peptide gene exhibit enhanced resistance against *Xvm*, providing a strong foundation for engineering durable resistance to BXW. The synergistic stacking of *MusaVicilin* with other defense-related genes that could further strengthen and broaden resistance, offering a more robust defense strategy. Further evaluation of these transgenic events under confined field trials is essential to validate their disease resistance efficacy and agronomic performance.

## Author’s contribution

Concept conceived by LT; Experiments performed by SWM, and designed by JNT, VON, SMK. All authors reviewed the manuscript.

## Acknowledgements

The authors wish to thank International Institute of Tropical Agriculture (IITA) for funding support. The first author wishes to thank for the Ph.D. opportunity by IITA.

